# BiP/GRP78 is a pro-viral factor for diverse dsDNA viruses that promotes the survival and proliferation of cells upon KSHV infection

**DOI:** 10.1101/2023.09.29.560238

**Authors:** Guillermo Najarro, Kevin Brackett, Hunter Woosley, Catya Faeldonea, Osvaldo Kevin Moreno, Adriana Ramirez Negron, Christina Love, Ryan Ward, Charles Langelier, Brooke M. Gardner, Carolina Arias

## Abstract

The Endoplasmic Reticulum (ER)-resident HSP70 chaperone BiP (HSPA5) plays a crucial role in maintaining and restoring protein folding homeostasis in the ER. BiP’s function is often dysregulated in cancer and virus-infected cells, conferring pro-oncogenic and pro-viral advantages. We explored BiP’s functions during infection by the Kaposi’s sarcoma-associated herpesvirus (KSHV), an oncogenic gamma-herpesvirus associated with cancers of immunocompromised patients. Our findings reveal that BiP protein levels are upregulated in infected epithelial cells during the lytic phase of KSHV infection. This upregulation occurs independently of the unfolded protein response (UPR), a major signaling pathway that regulates BiP availability. Genetic and pharmacological inhibition of BiP halts KSHV viral replication and reduces the proliferation and survival of KSHV-infected cells. Notably, inhibition of BiP limits the spread of other alpha- and beta-herpesviruses and poxviruses with minimal toxicity for normal cells. Our work suggests that BiP is a potential target for developing broad-spectrum antiviral therapies against double-stranded DNA viruses and a promising candidate for therapeutic intervention in KSHV-related malignancies.

## Introduction

Viruses dramatically remodel cellular physiology to accommodate the heightened biosynthetic demand for generating new viral particles. This process is orchestrated by virus-encoded factors that subvert the host protein homeostasis (proteostasis) machinery to promote the timely and optimal synthesis, folding, and maturation of proteins required for viral replication ^1,2^. Amongst the hundreds of proteostasis factors, molecular chaperones are critical for viral infections ^2–5^.

Molecular chaperones assist in folding, refolding, and translocating nascent, unfolded, or misfolded proteins to promote the acquisition of functional conformations or target terminally misfolded proteins for degradation, thus maintaining proteome integrity ^6,7^. Viruses co-opt host chaperones, especially those belonging to the heat shock protein (HSP) family, by altering the levels, interactions, or localization of HSPs to facilitate viral entry and replication, viral protein synthesis, and virion assembly ^2,3^. While all viruses exploit the function of chaperones in different cellular compartments, enveloped viruses, which are surrounded by an outer lipid layer acquired from the host and encode one or more glycoproteins, heavily rely on endoplasmic reticulum (ER) chaperones ^5,8^.

The ER is a membrane-bound organelle where most transmembrane and secretory proteins are synthesized, folded, and modified ^9^. A master regulator of ER functions is the Binding immunoglobulin protein/Glucose-regulated protein 78 (BiP/GRP78), an ER-resident HSP70 that assists nascent peptide folding ^10^. BiP is also a key player in the unfolded protein response (UPR), the ER stress response ^11–13^. BiP modulates the activity of the three transmembrane ER stress sensor proteins governing the UPR. These sensors are the kinase/nuclease IRE1, the kinase PERK, and the ER-membrane tethered transcription factor ATF6 ^14,15^. When the cell’s biosynthetic output surpasses the ER’s folding capacity, unfolded proteins accumulate in the ER lumen, which licenses the activation of IRE1, PERK, and ATF6 through well-described mechanisms involving their reversible dissociation from BiP, direct activation by unfolded protein ligands, or changes in the redox status of the ER lumen. The signaling cascade downstream of the sensors culminates in the activation of gene expression programs that restore ER homeostasis (reviewed in ^16^).

In virus-infected cells, BiP is upregulated by flaviviruses (Zika, ZIKV; and Dengue, DENV virus), coronaviruses (SARS-CoV-2, MERS-CoV), Hepatitis B Virus (HBV), and Human Cytomegalovirus (HCMV), a member of the betaherpesvirus subfamily ^17–21^. Pharmacological inhibition or knockdown of BiP reduces viral replication in cultured cells (ZIKV, DENV, SARS-CoV-2, and HCMV) and mouse infection models (SARS-CoV-2), highlighting its pro-viral activity.

Beyond viral infection, BiP is elevated in numerous cancers, including leukemia, melanoma, multiple myeloma, brain, pancreatic, liver, and breast cancer, and is regarded as a promising biomarker and therapeutic target in several diseases ^22^. In addition to heightened levels, BiP can re-localize to the cell surface during ER stress, which correlates with tumor aggressiveness and poor prognosis ^23,24^. Moreover, BiP protects cancer cells from apoptosis and promotes proliferation and metastasis, thereby contributing to tumor robustness and resistance to therapy _22,25._

Motivated by these observations, we investigated the roles of BiP during infection by the oncogenic gamma-herpesvirus Kaposi’s Sarcoma-Associated Herpesvirus (KSHV). KSHV is the most recently discovered human herpesvirus and the causal agent of Kaposi’s sarcoma (KS), the lymphoproliferative disorders Primary Effusion Lymphoma (PEL), and KSHV-associated multicentric Castleman’s disease (MCD), and is implicated in KSHV inflammatory cytokine syndrome (KICS) ^26,27^. Few treatment options are available for these diseases, and an unmet clinical need exists for a targeted antiviral therapeutic ^27^.

KSHV contains a ∼160Kb double-stranded DNA genome that encodes over 80 proteins ^28,29^. As with all other herpesviruses, KSHV establishes life-long latent infections characterized by the expression of a few viral products. To complete its life cycle, KSHV reactivates from this latent state to a lytic, virion-productive infection characterized by a massive induction of viral transcripts and protein synthesis ^29,30^. KSHV is an enveloped virus, and several of its proteins are synthesized in the ER ^31^; therefore, KSHV infection could impose a high biosynthetic burden on this organelle.

Here, we show that BiP is upregulated during KSHV lytic infection, independent of UPR activation, and acts as a pro-viral factor for multiple types of DNA viruses (herpes and poxviruses), underscoring that inhibiting BiP may provide broad-spectrum antiviral utility. Moreover, we report that BiP inhibition with the thiazole benzenesulfonamide HA15 has strong cytostatic and cytotoxic effects in KSHV-infected B-cells and primary endothelial cells but not in uninfected cells, supporting the notion that BiP inhibition is a promising therapeutic alternative for KSHV-associated malignancies.

## Results

### BiP is upregulated during the lytic cycle of KSHV

KSHV encodes at least 14 transmembrane or secreted proteins with functions in cell entry, viral gene expression, and immune evasion (Table 1). These viral proteins are folded, processed, and assembled in the ER with the assistance of cellular chaperones. To investigate the role of BiP during the lytic cycle of KSHV, we used the well-established iSLK.219 model system to study KSHV reactivation ^32^. iSLK.219 cells are latently infected with KSHV and contain a doxycycline (Dox)-inducible viral transcription factor RTA (replication and transcriptional <u>a</u>ctivator), the expression of which is sufficient to induce entry to the KSHV’s lytic cycle (Fig. 1A). iSLK.219s harbor KSHV.219, a recombinant virus that encodes a constitutive GFP reporter and an RTA-inducible RFP reporter in the viral genome that facilitates monitoring infection and viral reactivation (Sup. Fig. 1A-B)^33^. We induced iSLK.219 cells with Dox and collected cell lysates at 0h, 24h, 48h, and 72h, representing the latent (0h), early-lytic (24h-48h), and late-lytic (48h-72h) stages of infection, and monitored the levels of BiP by immunoblot throughout a time course of reactivation (Fig. 1B). Protein levels of BiP significantly increased early in the lytic cycle of KSHV, starting at 24h post-reactivation, and coincide with an upsurge in viral protein expression (Fig. 1B and Sup. Fig. 1B). To determine the timing of BiP upregulation during the lytic cycle, we used the viral DNA replication inhibitor phosphonoformate (PFA), which arrests infection in the early stages of the lytic cycle by preventing viral DNA replication ^34^. The levels of BiP in iSLK.219 cells induced with Dox for 72h were indistinguishable in PFA-treated from untreated cells, indicating that the upregulation of BiP is an early event in the viral lytic cycle that is independent of late viral gene expression (Fig. 1C). Notably, we did not detect any changes in the levels of GRP94 or Calreticulin—two prominent ER chaperones—during the lytic cycle of KSHV, suggesting that the upregulation of BiP during infection is not general to all ER chaperones (Fig. 1D).

**Figure 1.**
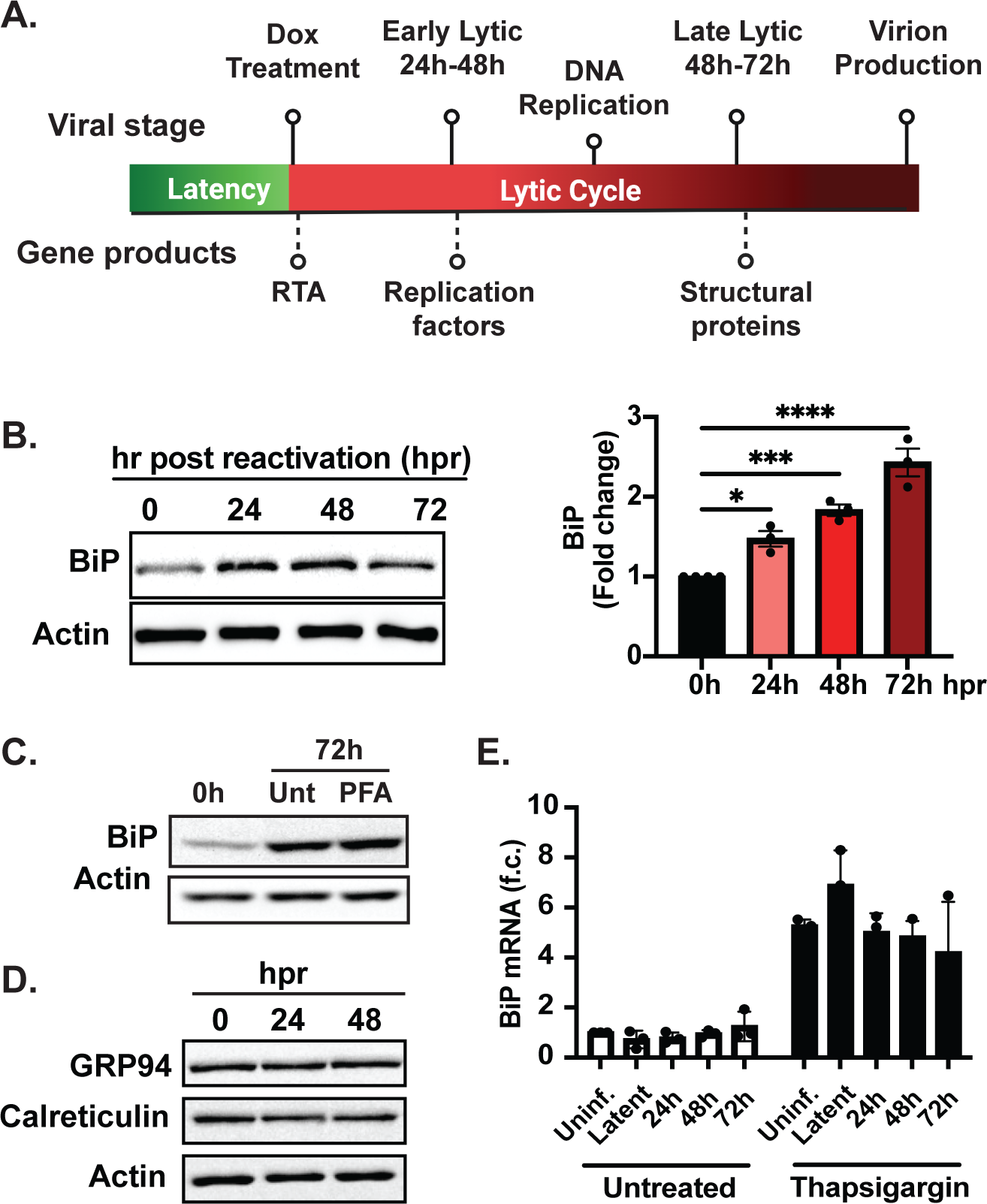
BiP is upregulated during the KSHV lytic cycle. **(A.)** Schematic of lytic reactivation in iSLK.219 cells **(B.)** BiP is upregulated at the protein level in a time course of reactivation. (Left) iSLK.219 cells were treated with Dox (1µg/ml) to induce RTA expression and viral reactivation. Whole-cell lysates collected at the indicated times were analyzed by immunoblot. Actin: loading control. (Right) Image densitometry quantification of the immunoblot **(C.)** BiP upregulation is independent of late viral gene expression. Viral DNA replication was inhibited in iSLK.219 cells by pretreatment with PFA (100nM) for 24 hours before induction with Dox. Whole-cell lysates collected at the indicated times were analyzed by immunoblot. Actin: loading control. **(D.)** Immunoblot of GRP94, calreticulin, and actin during KSHV reactivation in iSLK.219 cells **(E.)** BiP upregulation is post-transcriptional. qRT-PCR quantification of BiP mRNA in a time course of reactivation in iSLK.219 untreated or treated with Tg (100nM) for 4h. Note the high levels of BiP mRNA in cells undergoing acute ER stress. *N*=3 independent biological replicates for (B, C, D, and E). Values in (B, E) are average ±SEM. Statistical significance in (B) was calculated using a one-way ANOVA (*P=0.01, **P=0.004, **P=<0.0001).

**Table 1:** KSHV proteins containing signal peptides. The protein sequences of all annotated KSHV proteins (GQ994935.1) were analyzed with the signal sequence prediction engines Phobius^69^, Signal P 6.0^70^, and Predisi^71^. Signal peptides (SP) were annotated if predicted by two or more engines. The transmembrane domains of proteins containing SPs were annotated using DeepTMHMM^72^.

| Gene Name | Function | Time of Expression | Signal Peptide | Transmembrane domains |
| --- | --- | --- | --- | --- |
| K1 | Glycoprotein | Latent | 1-18 | 224-247 |
| ORF4 | Complement Binding Protein | Early | 1-19 | 533-551 |
| ORF8 | Glycoprotein B | Late | 1-26 | 743-762 |
| K2 | Viral Interleukin 6 Homologue | Latent | 1-22 | N/A |
| K4 | v-Macrophage Inflammatory Protein 2 | Immediate Early | 1-25 | N/A |
| K4.1 | v-Macrophage Inflammatory Protein 3 | Immediate Early | 1-27 | N/A |
| K6 | v-Macrophage Inflammatory Protein 1 | Immediate Early | 1-24 | N/A |
| ORF22 | Glycoprotein H | Late | 1-21 | 715-736 |
| ORF39 | Glycoprotein M | Early | 1-29* | 14-25, 79-103, 119-135, 153-171, 211-231, 240-260, 274-293, 307-325 |
| ORF47 | Glycoprotein L | Early | 1-20 | N/A |
| K8.1 | Glycoprotein | Late | 1-26 | 200-220 |
| ORF53 | Glycoprotein N | Late | 1-23 | 79-99 |
| K14 | Viral OX2 | Early | 1-24 | 230-250 |
| K15 | LMP1/2 Homologue | Latent/Early | 1-25 | 10-25, 35-50, 69-81, 91-100, 123-139, 150-167, 178-194, 207-223, 240-250, 272-287, 299-308, 329-349 |

### BiP upregulation during the early-lytic cycle of KSHV is independent of the UPR

BiP transcription is rapidly upregulated by the UPR transcription factors XBP1s and ATF6 in response to ER stress to allow homeostatic readjustment ^13,35^. qRT-PCR analysis revealed that KSHV lytic infection did not coincide with an increase in BiP mRNA levels (Fig. 1E), suggesting that the upregulation of the BiP protein we observed was post-transcriptional and likely independent of UPR induction during infection. Previous reports indicate that KSHV modulates the UPR in PEL-derived cells by disrupting the signal transduction downstream of the UPR sensors IRE1, PERK, and ATF6 ^36^. To determine whether the UPR is dysregulated during KSHV infection in iSLK.219 cells, we first evaluated the levels, phosphorylation status, and activity of IRE1 during viral reactivation (Sup. Fig. 2A). Phosphorylated IRE1 (IRE1-P) levels increased as the lytic cycle progressed. Despite the evident activation of IRE1 during the lytic cycle of KSHV, we observed a minimal increase in XBP1 mRNA splicing (XBP1s) and XBP1s protein, a direct product of IRE1 activity, indicating disruption of canonical IRE1 signaling during lytic infection in iSLK.219 cells (Sup. Fig. 2A-C).

Even though XBP1s was barely detectable during the KSHV lytic cycle in iSLK.219 cells, we tested whether the low levels of this potent UPR transcription factor could mediate the upregulation of BiP. We used CRISPRi-mediated gene silencing of XBP1 to test this hypothesis (Sup. Fig. 2D) ^37^ and found that the knockdown of XBP1 did not significantly impact BiP protein levels or viral production in iSLK.219 cells (Sup. Fig. 2D, 2E). We also investigated whether ATF6 was responsible for the upregulation of BiP observed during KSHV lytic infection. To this end, we knocked down ATF6 or treated iSLK.219 cells with the ATF6 inhibitor CeapinA7, a small molecule that blocks ATF6 ER export and its subsequent proteolytic processing and activation ^38^. Neither CeapinA7 treatment nor ATF6 knockdown by CRISPRi affected the accumulation of BiP protein or the production of infectious virus in iSLK.219 cells undergoing KSHV lytic infection (Sup. Fig. 2F-H). These findings establish that KSHV reactivation exerts UPR-independent, post-transcriptional BiP protein upregulation in these cells (Sup. Fig. 2D, 2F-G).

### BiP is a pro-viral factor in KSHV-infected cells

The upregulation of BiP protein during KSHV lytic infection in iSLK.219 cells is remarkable given the substantial host shutoff mediated by the KSHV SOX (shutoff and exonuclease) protein, which degrades host mRNAs. This viral factor suppresses the expression of host proteins to funnel host cell resources towards the pathogen’s benefit ^39^. Because this result suggests BiP effectively escapes host shutoff in iSLK.219 cells, we tested whether KSHV exploits BiP during its lytic cycle. To this end, we used HA15, a thiazole benzenesulfonamide inhibitor of BiP that targets its ATPase domain ^40^. Treatment of iSLK.219 cells with HA15 had a striking effect on KSHV reactivation by reducing lytic protein expression and decreasing infectious virus production by up to 90% (Fig. 2A-B). In an orthogonal approach, we silenced BiP expression by siRNA-mediated knockdown. BiP’s genetic depletion phenocopied HA15 treatment and significantly reduced viral protein expression and infectious virus production (Fig. 2C-D), thus corroborating that BiP is essential for KSHV replication in these cells. To determine if the observed effect of HA15 was restricted to iSLK.219 cells, we investigated its impact on the inducible B-cell lymphoma-derived cell line TREx-BCBL1-RTA, which is also latently infected with KSHV and expresses RTA under the control of a doxycycline-inducible promoter ^41^. Interestingly, in TREx-BCBL-1 cells, we did not detect an upsurge in BiP protein levels during the KSHV lytic cycle (Sup. Fig. 3). Despite this observation, treatment of TREx-BCBL-1-RTA cells with HA15 during a time course of lytic reactivation with Dox reduced viral protein expression and viral DNA replication comparable to the effect observed in iSLK.219s treated with HA15 (Fig. 2E-F). These observations confirm that BiP is a pro-viral factor during KSHV infection in multiple infection models and cell types.

**Figure 2.**
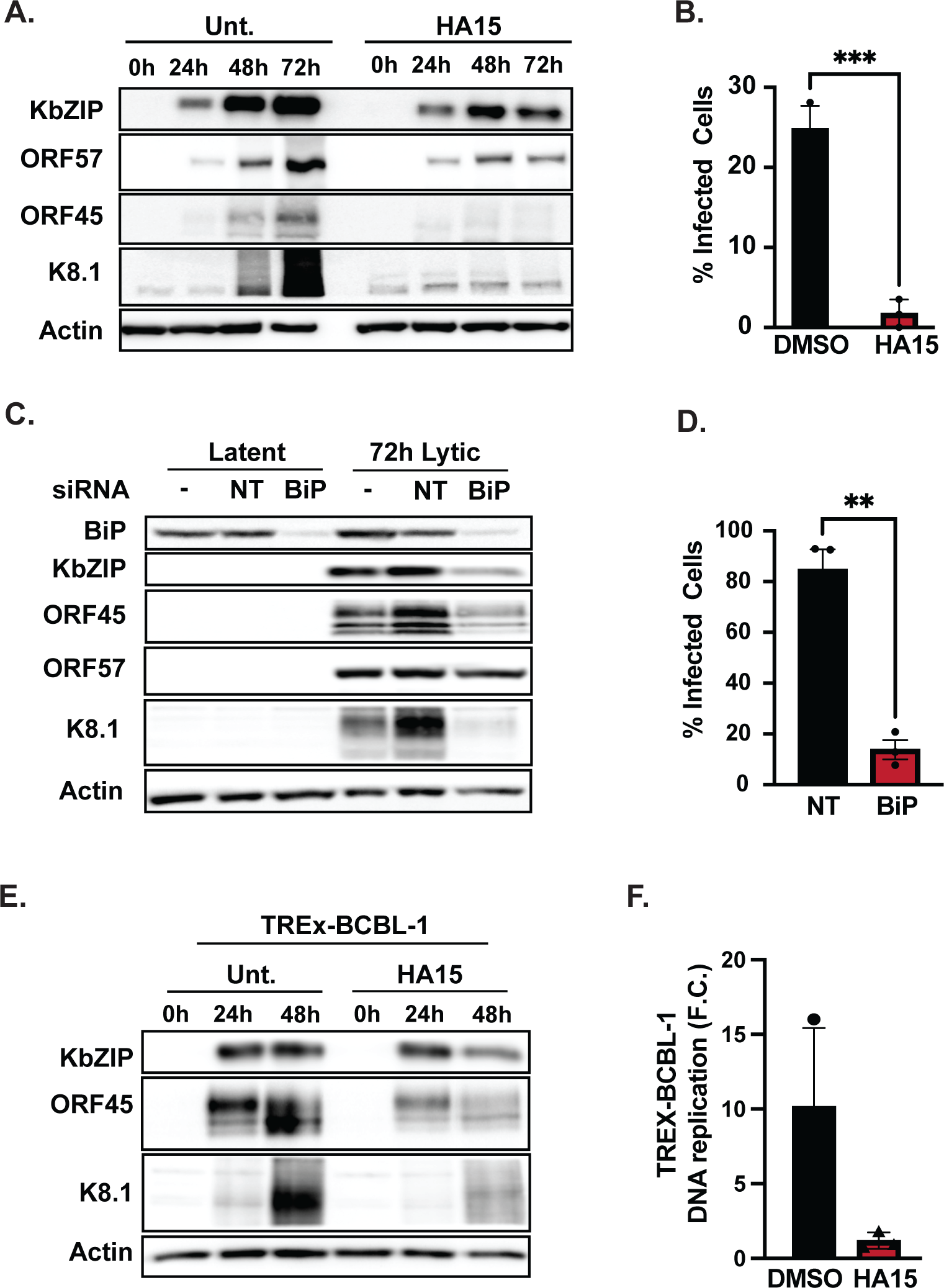
BiP is a pro-viral factor in KSHV-infected cells. **(A-B)** BiP inhibition with HA15 disrupts the lytic cycle iSLK.219s. **(A)** Cells were treated with HA15 (10µM) 24h before reactivation with Dox (1µg/ml). At the indicated times, whole-cell lysates were collected and analyzed by immunoblot for viral proteins (Immediate early: KbZip-nuclear, ORF57-nuclear, Early: ORF45-nuclear/cytosolic, Late: K8.1-glycoprotein). Actin: loading control. **(B)** Supernatants from iSLK.219 cells treated with HA15 were collected at 72h post reactivation and used to infect naïve iSLK cells. GFP expression was determined by automated cell counting at 48h post-infection and used as a proxy for virus production. **(C-D)** Silencing of BiP reduces viral reactivation and infectious virion production. **(C)** iSLK.219 cells were reactivated with Dox, following BiP siRNA-mediated silencing for 48h. Lysates were collected at 72h post-reactivation and analyzed by immunoblot for viral factors. siRNA – untransfected, NT non-targeting **(D)** Supernatants from BiP-KD cells treated as in (C) were collected and processed as described in (B). **(E-F)** Inhibition of BiP blocks the lytic cycle in TREx-BCBL-1-RTA cells. **(E)** Cells were treated with HA15 (10µM) for 24h before induction with Dox (1µg/ml). At 48h post-infection, whole cell lysates were collected and analyzed by immunoblot. Actin: loading control. **(F)** Total DNA was isolated from cells treated as in (E), and viral DNA was quantified by qRT-PCR. *N*=3 independent biological replicates. Values in (B, D, F) are average ±SEM. Statistical significance was calculated using a paired t-test (B and D) (*P=0.01, **P=0.002) or a one-way ANOVA (F) (** P=0.0065)

### BiP inhibition disrupts the early stages of the KSHV lytic cycle

Our observations suggested that blocking BiP function disrupts the KSHV lytic cycle at early stages post reactivation. To test this hypothesis, we analyzed viral transcriptomes collected by RNAseq at 72h post reactivation to determine the impact of BiP inhibition/depletion on viral gene expression at a genome-wide level (Fig. 3A-B). As anticipated, we observed a global reduction in viral transcript levels during the lytic cycle of HA15-treated cells, except for the K2 transcript (encoding vIL6, the viral homolog of interleukin 6) and its overlapping transcript ORF2 (encoding a viral dihydrofolate reductase), both of which increase at 72h post reactivation in HA15-treated cells compared to untreated cells (Fig. 3A) ^29,42^. Previous reports have found XBP1s can bind to the promoter of vIL6 in KSHV-infected cells to induce its expression ^43^. Considering that BiP inhibition by HA15 can cause ER stress and UPR activation, we measured the protein levels of XBP1s in a time course of reactivation in the presence of HA15. In these conditions, we could not detect the expression of XBP1s protein in HA15-treated iSLK.219 cells, suggesting that additional factors may compensate for upregulating vIL6 (Sup. Fig. 4).

**Figure 3.**
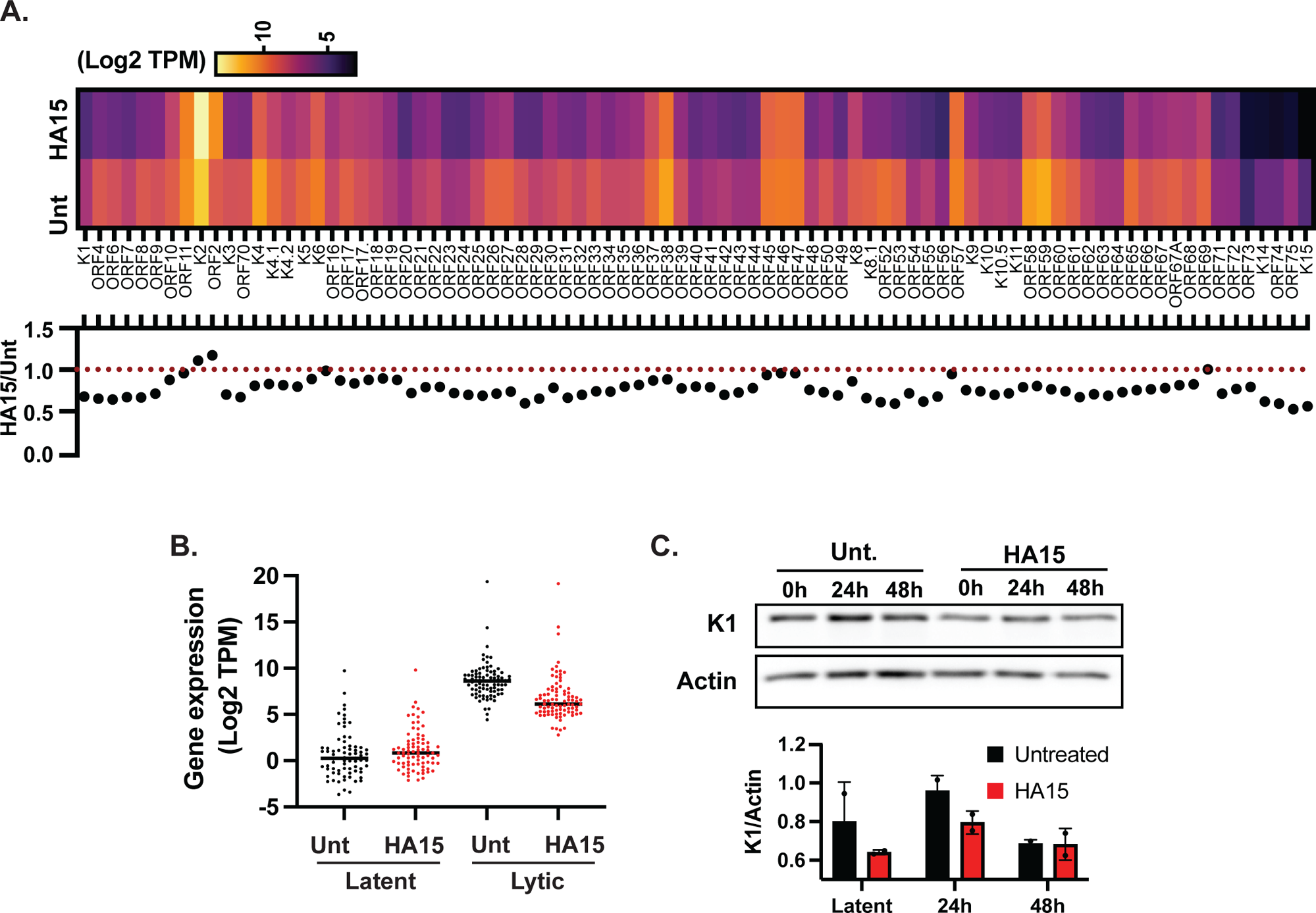
BiP inhibition disrupts the KSHV lytic cycle. **(A-B).** Total RNA was isolated from latent and lytic iSLK.219 cells in the presence or absence of HA15. RNAseq libraries were prepared, sequenced, and aligned to the KSHV genome. **(A.)** (Top) Heatmap showing the Log2 of transcripts per million (TPM) for all KSHV genes ordered by genomic position in lytic iSLK.219 cells ± HA15. (Bottom) Ratio of Log2 of TPM in HA15 vs. untreated (Unt). The dotted red line at ratio=1 denotes no change in viral gene expression in HA15-treated cells vs. untreated cells **(B.)** Boxplot of the Log2 of TPM of KSHV genes in latent and lytic iSLK.219 cells at 72h post-reactivation in the presence or absence of HA15. **(C.)** HA15 treatment reduces K1 levels during the KSHV lytic cycle. (top) TREx-BCBL-1-RTA cells were treated with HA15 (10µM) 24h before induction with Dox (1ug/ml). At 48h post-infection, whole cell lysates were collected and analyzed by immunoblot. Actin: loading control. (bottom) Image quantification by gel densitometry of the K1 immunoblot. *N=* 3 independent biological replicates. Values in (C) are average ±SD.

Given the essential role of BiP for folding and processing newly synthesized proteins in the ER, we hypothesized that HA15 treatment could disrupt the lytic cycle by affecting the function of viral glycoproteins expressed early after reactivation (Table 1). We focused on K1, a KSHV glycoprotein expressed during the latent and lytic cycles of infection, which is required for efficient lytic reactivation ^44,45^. We evaluated the levels of K1 by immunoblot in TREx-BCBL-1-RTA cells and observed an increase in the levels of this protein as the lytic cycle progressed, in agreement with previous findings ^45^ (Fig. 3C). We see that the levels of K1 are generally lower in latent and lytic cells treated with HA15, which may negatively impact the progress of the lytic cycle thus suggesting that BiP inhibition could disrupt the KSHV lytic cycle in part by modulating K1 levels.

### HA15 is a broad-spectrum inhibitor of herpes- and poxvirus replication

BiP inhibition with HA15 has been shown as a potential antiviral strategy for RNA viruses, including alphaviruses and, more recently, coronaviruses ^19,46^. Our results indicate that this compound is also active against KSHV. These observations raised the possibility that HA15 may provide antiviral utility against other dsDNA viruses. To test this hypothesis, we evaluated the potential of HA15 to inhibit viral replication in primary human fibroblasts (NHDFs) infected with three different dsDNA viruses: an alphaherpesvirus, Herpes Simplex Virus-1 (HSV-1), a betaherpesvirus, Human Cytomegalovirus (HCMV), and a poxvirus, Vaccinia Virus (VV). Cells were infected at a low multiplicity of infection (MOI) in the presence or absence of HA15. The spread of infection at different times post-infection was determined by measuring the expression of virus-encoded GFP in HSV-1-GFP and HCMV-GFP infected cells or by immunofluorescence using a polyclonal antibody against vaccinia virus ^47–49^. Our experiments revealed potent inhibition of viral spread for HSV-1-GFP, HCMV-GFP, and VV in the presence of 10-30 µM HA15, indicating that HA15 acts as a broad-spectrum inhibitor of dsDNA viruses (Fig. 4A). Notably, HA15 treatment of NHDFs was not cytotoxic even at high concentrations (30 µM) or long treatment times (1 or 6 days) (Fig. 4B), further substantiating that blocking BiP is a promising antiviral strategy with a minimal negative impact on normal cells.

**Figure 4.**
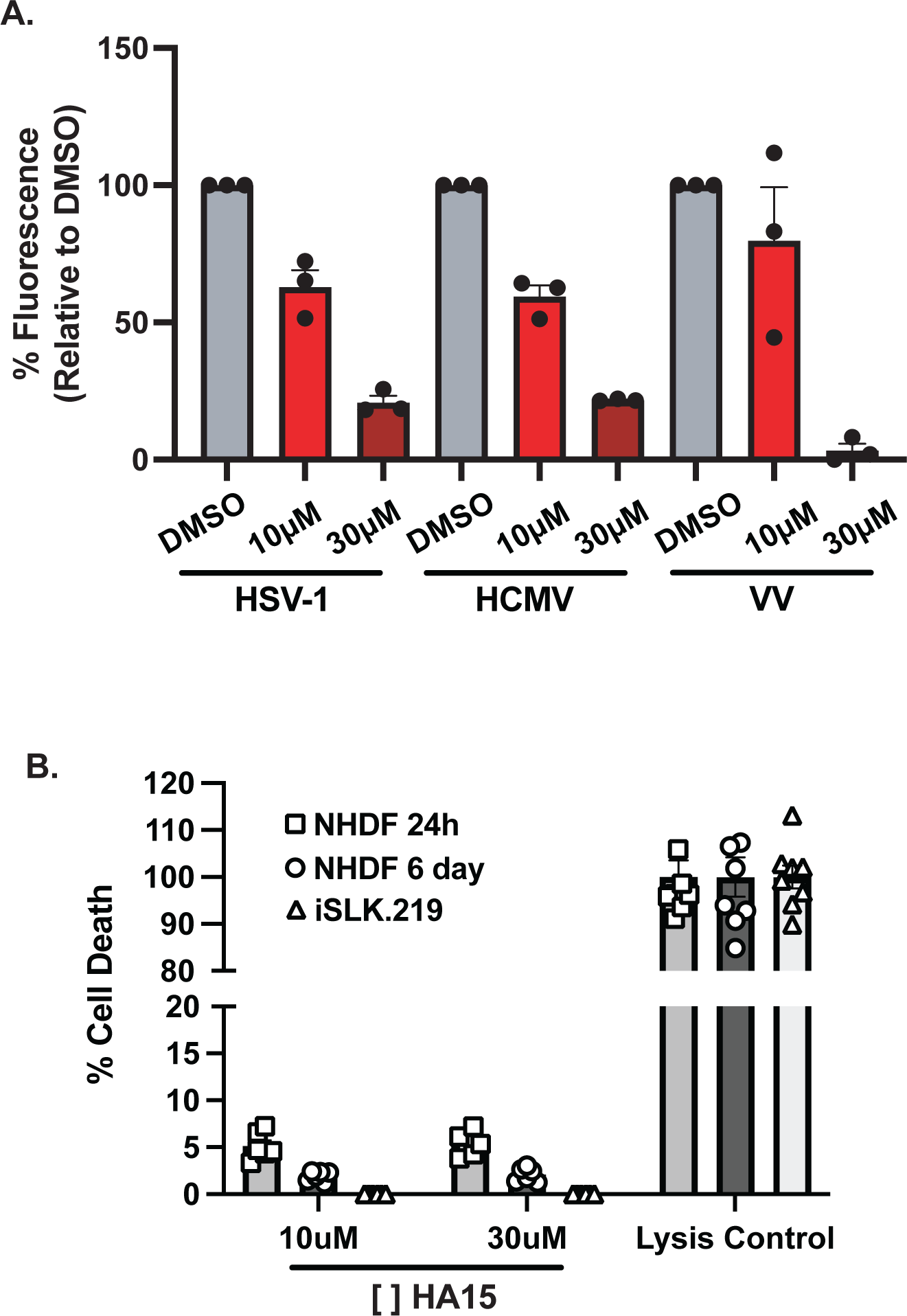
The BiP inhibitor HA15 has a broad-spectrum antiviral effect on herpesviruses and poxviruses. **(A.)** Primary human fibroblasts (NHDF) were Infected at a low multiplicity of infection (MOI) (HSV-1 MOI 0.001, HCMV at MOI 0.1, and VV at MOI 0.01 in the presence or absence of HA15 (10uM or 30uM). The spread of infection was determined at different times post-infection by measuring the expression of virus-encoded GFP in HSV-1-GFP and HCMV-GFP infected cells or by immunofluorescence using a polyclonal antibody against Vaccinia virus. **(B)** The effect of HA15 treatment on the viability of NDHF (1 or 6 days) and iSLK.219s (3 days) was evaluated by measuring LDH release. N=3 (A), N=6 (B) independent biological replicates. Values are average ±SEM.

### Treatment with HA15 is cytostatic for KSHV-infected lymphoma-derived B-cells

In addition to its antiviral activity, HA15 has shown promising anticancer activity ^40,50^. To test whether this compound has a similar anticancer effect in KSHV-related lymphomas, we evaluated the impact of escalating doses of HA15 on the viability of three cell lines derived from primary effusion lymphoma, TREx-BCBL-1-RTA, and the BC-1 and BC-2 cell lines that are co-infected with KSHV and EBV (Epstein Barr Virus) (Fig. 5A-B, Sup. Fig. 5A-D). At 72h post-treatment, we observed a dose-dependent reduction in cell numbers for these cancer cell lines. Even as the total cell numbers were lower in HA15 treatment, the viability of treated cells remained essentially unchanged in all three cell lines at HA15 concentrations ≤10 µM. The highest HA15 concentration we tested (50µM) resulted in profound cell cytotoxicity measured by trypan blue exclusion (Fig. 5A-B, Sup. Fig. 5A-D). These observations suggest HA15 (1-10µM) has a strong cytostatic effect in B-cells derived from primary effusion lymphoma and is cytotoxic to cancer cells at high concentrations. Finally, to test whether these HA15 effects are specific to cancer cells, we treated non-transformed normal peripheral primary B cells (PPBCs) with increasing doses of HA15. These experiments revealed no significant changes in the total number of viable cells compared to untreated cells, even at the highest concentration tested (50µM), indicating that HA15 is neither cytostatic nor cytotoxic for normal B-cells (Fig. 5C-D).

**Figure 5.**
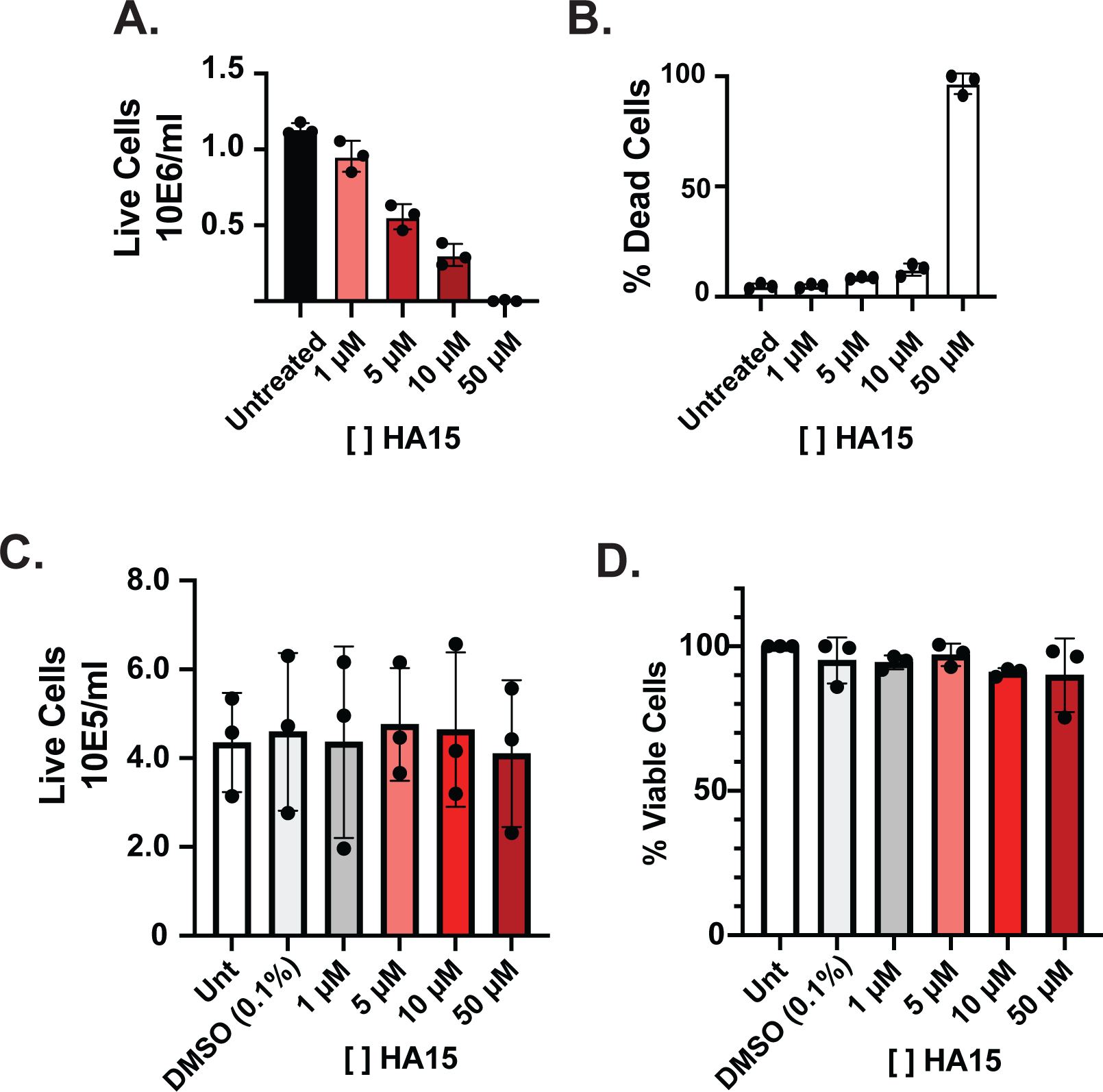
BiP Inhibition with HA15 causes strong cytostasis in latent PEL-derived cells. **(A-B.)** HA15 treatment differentially reduces cell numbers compared to cell viability in TREx-BCBL-1 cells. Latent TREx-BCBL-1 cells were treated with increasing concentrations of HA15 (0-50 μM) for 72 hrs. The total number of viable cells **(A.)** and the percent of dead TREx-BCBL-1 cells **(B.)** were determined by automated cell counting following trypan blue staining. **(C-D.)** HA15 treatment does not cause cytostasis nor cytotoxicity in primary B cells. Primary Peripheral B-cells were treated with increasing concentrations of HA15 (0-50 μM) for 72 Hrs. The total number of viable cells **(C.)** and the percent of live cells **(D.)** were determined as described in (A-B). *N*=3 independent biological replicates. Values are average ±SEM.

### Treatment with HA15 is cytotoxic for KSHV-infected primary lymphatic endothelial cells

The main cellular targets of KSHV in KS lesions are spindle cells thought to originate from lymphatic endothelial cells (LECs) ^51,52^. We used primary LECs as a model for KSHV infection to study the effects of BiP inhibition in a context relevant to the pathophysiology of KS. In this model, we infected LECs with the recombinant KSHV.219 virus, which harbors a puromycin resistance cassette, a constitutive GFP reporter, and an RTA inducible RFP reporter ^33,53^. At 14 days post-infection and following puromycin selection (started 48h after infection), KSHV-infected LECs (KLECs.219) expressed GFP and showed the typical spindle cell morphology that is characteristic of KS lesions, corroborating KSHV infection (Fig. 6C top middle and left panels). As previously reported, a small fraction of KLECs.219 expressed RFP (data not shown), indicating spontaneous lytic reactivation in cell culture ^53^. In line with our findings in iSLK.219 cells, we observed the upregulation of BiP in KLECs.219 at 14 days post-infection, possibly driven by the expression of lytic genes in a subset of the population (Fig. 6A). Treatment of uninfected LECs with 10 µM HA15 for up to 72 h did not substantially affect cell morphology or viability (Fig. 6B-C). Remarkably and in stark contrast to uninfected LECs, treating KLECs.219 with 10µM HA15 for 72h induced significant cell death (Fig. 6B-C), with evident cytotoxicity as early as 48h post-treatment.

**Figure 6.**
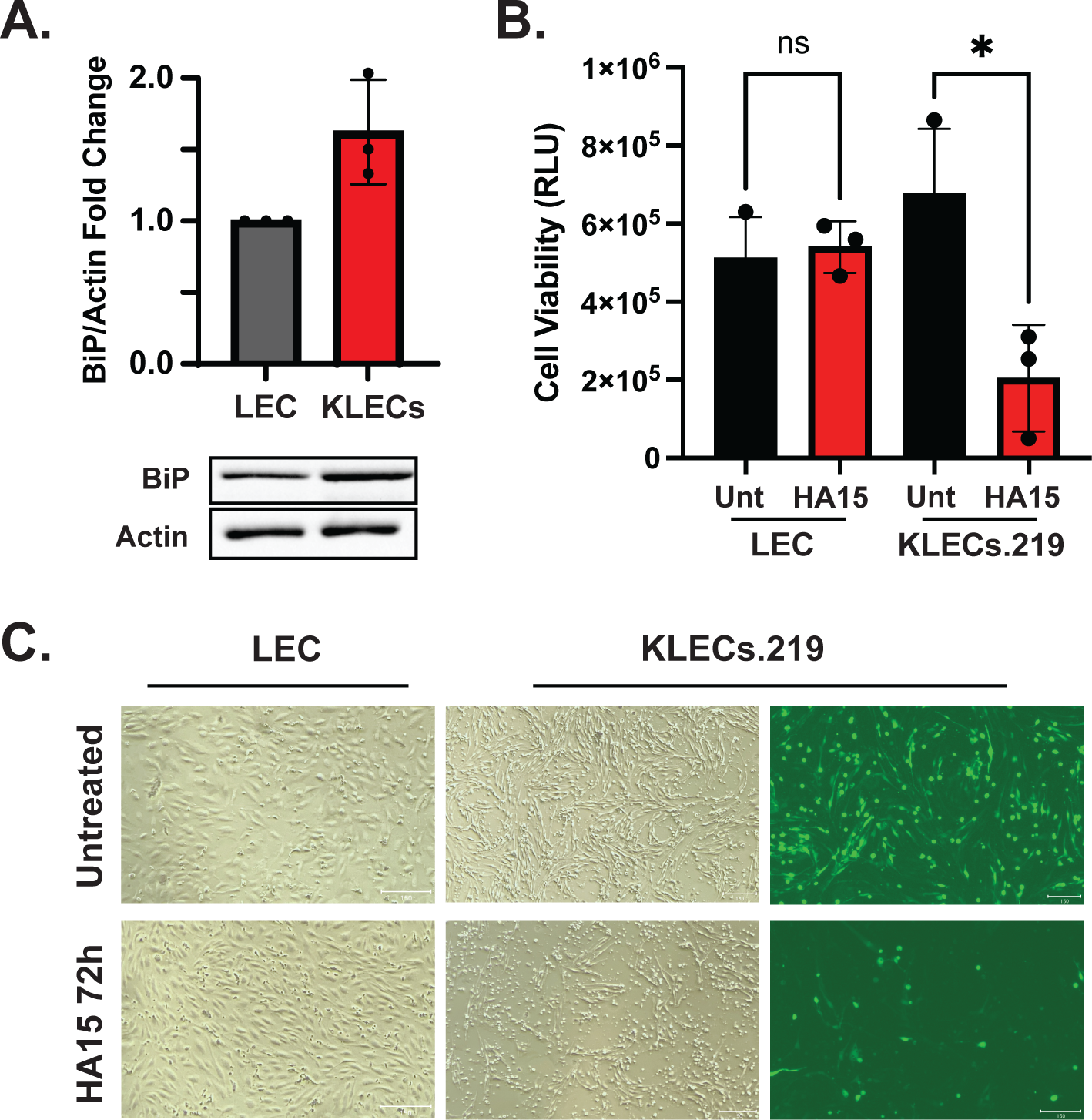
HA15 has a cytotoxic effect on KSHV-infected LEC cells. Primary Lymphatic endothelial cells were infected with KSHV.219 and selected with puromycin for 7-14 days. **(A.)** Whole-cell lysates from uninfected (LEC) or infected (KLECs) were collected and analyzed by immunoblot. Actin: Loading control. **(B-C)** LECs and KLECs were treated with HA15 (10uM) for 72h. Cell viability was evaluated by ATP quantification using CellTiter-Glo **(B.)** and microscopy at 0h and 72h post-treatment **(C.)**.

## Discussion

BiP is a pivotal component of the proteostasis network and a pro-viral factor; therefore, it emerges as a potential target for antiviral intervention. Our study uncovered the dysregulation and requirement for BiP during lytic infection by the oncogenic herpesvirus KSHV. Furthermore, we showed that the BiP inhibitor HA15 had a broad-spectrum antiviral activity for dsDNA viruses (herpesviruses and poxviruses) and caused cytostasis/cytotoxicity in KSHV-infected PEL and LEC cells, highlighting its potential use as an anticancer agent during viral-induced oncogenesis.

The cellular response to ER proteostatic insults is orchestrated by the UPR, wherein BiP upregulation is mainly transcriptionally driven by the UPR transcription factors XBP1s and ATF6 ^54^. In KSHV-infected iSLK.219 cells, BiP escaped UPR regulatory control and was upregulated post-transcriptionally during lytic infection. Viral infections, including KSHV, induce the integrated stress response (ISR), which has, as a principal outcome, the downregulation of global protein synthesis ^55^. In these conditions, cap-dependent translation is disfavored. Thus, the enhanced BiP protein synthesis we observed may arise from alternative initiation mechanisms such as the one afforded by the IRES element in the BiP mRNA ^56,57^. Indeed, several stresses negatively impact cap-dependent translation to favor the expression of IRES-containing transcripts ^58^. Another possibility is that the translation initiation factor eIF2A, not to be confused with eIF2alpha, promotes the translation of BiP during KSHV infection, as reported in other stress conditions ^59^. In this scenario, eIF2A would facilitate translation initiation at non-canonical start codons in situations in which eIF2alpha is phosphorylated (i.e., during ISR activation), and the availability of the eIF2-Met-tRNAi-GTP ternary complex, which is required for cap-dependent translation initiation at cognate AUG start codons, is limiting. Whether these mechanisms—IRES- or eIF2A-mediated translation initiation—promote the BiP protein upregulation we observed remains to be determined. Nonetheless, the enhanced translation of upstream open reading frames and non-canonical start codons reported during the KSHV lytic cycle suggests an altered translational state that could account for the molecular phenotypes we observed ^29^.

The upregulation of BiP and its pro-viral activity extend beyond KSHV-inherent biology. Indeed, both BiP upregulation and pro-viral roles have been reported in corona-, flavi-, alphaherpes-, and betaherpesviruses ^11,19,46,60,61^. All these viruses are enveloped; therefore, they rely on the host’s machinery to acquire membranes and synthesize and correctly fold viral glycoproteins or secreted viral peptides^2^. In all the viruses mentioned above, BiP has been shown to participate in several steps in the viral cycle, attesting to its essential role in aiding the correct biosynthesis and assembly of proteins during virion production.

In cells infected with the alphavirus VEEV, the flaviviruses DENV or JEV, and the herpesvirus HCMV, genetically or pharmacologically blocking BiP does not impact viral genome replication but significantly reduces infectious virion production ^11,46,61^. In these cases, BiP may not be required for the early stages of the viral life cycle but for virion assembly. However, unlike the above observations on HCMV, VEEV, DENV, and JEV-infected cells, in KSHV-infected iSLK.219 and TREx-BCBL-1 cells, BiP inhibition results in a blockage of infection at early stages during reactivation of the lytic cycle before genome replication. This notion is substantiated by the lower levels of the early-lytic K1 viral glycoprotein we observed in lytic TREx-BCBCl-1 cells treated with HA15. Studies by the Damania Lab have shown that the expression of K1 is required for efficient KSHV replication ^45^, which is further supported by the global downregulation of viral gene expression, measured by RNASeq, that we observed in HA15-treated cells. Whether additional early-lytic proteins or host factors contribute to the downregulation of lytic reactivation in cells where BiP is no longer active remains to be determined.

While BiP was required for the efficient replication of KSHV in iSLK.219 epithelial cells and PEL-derived TREx-BCBL-1 cells, we noted that the levels of BiP did not increase in PEL-derived cells during the lytic cycle. The virus strains present in these cell lines (iSLK.219 Accession number GQ994935.1 and TREx-BCBL-1 Accession number HQ404500.1) are greater than 99% similar at the nucleotide sequence level (data not shown), suggesting that the disparate responses we observed likely stem from cell-intrinsic factors. PEL-derived cells show a gene expression profile resembling malignant plasma cells, including a higher expression of the UPR effector XBP1s, and, indeed, higher levels of XBP1s have been observed during the KSHV lytic cycle in TREx-BCBL-1 cells than those observed in iSLK.219 cells ^36,62^. The unique gene expression profile of PEL-derived cells may indicate profound reconfiguration of the machinery required for maintaining ER homeostasis in cells with a high secretory burden, as occurs in plasma cells ^62,63^. As such, in TREx-BCBL-1 cells the capacity of the ER may be sufficient to accommodate KSHV protein folding during the lytic cycle without a need to induce signal transduction programs to increase BiP levels. Future studies comparing the basal levels of BiP and other UPR factors in KSHV-infected B- and epithelial cells, as well as the identity and dynamics of BiP client proteins during the viral lytic cycle, will shed light on the inherent ER-protein folding capacity of different KSHV-infected cell types.

Our observations align with the cytoprotective role of BiP, particularly under stress conditions. In line with a maladaptive dependency on BiP in cancer cells, blocking BiP in KSHV-infected PEL and LEC-derived cells resulted in cytostatic and cytotoxic responses, respectively. Indeed, BiP levels are associated with cell division and increased proliferation rates in numerous tumor models ^22,64^. One mechanism by which BiP may confer a maladaptive survival advantage is through modulation of cell proliferation by tuning Wnt/B-catenin signaling, wherein BiP-Wnt interactions promote Wnt’s correct posttranslational processing to promote downstream signaling ^65^. In PEL cells, the Wnt/B-catenin signaling pathway is usurped by KSHV, and the latency-associated nuclear antigen (LANA), expressed in all KSHV-latently infected cells, arrests GSK3 in the nucleus and promotes the stabilization and accumulation of B-catenin, enabling the entry of infected cells into S-phase ^66^. Future experiments to evaluate the integrity of the Wnt/B-catenin signaling pathway in HA15-treated PEL cells will help clarify the contributions of BiP to changes in the proliferation capacity of these lymphoma-derived cells.

In contrast to PEL-derived cells, viral infection in KLECs.219 cells led to the upregulation of BiP and the strict dependence on BiP for cell survival. In other cancers, BiP inhibition leads to a hyperactive UPR, activating apoptosis and autophagy. Detailed mapping of host gene expression and proteome profiles following KSHV infection of LECs and treatment with HA15 will help determine which factors induced upon infection drive terminal responses in KLECs.219 cells.

One of the most exciting observations from our studies is the broad-spectrum antiviral activity of HA15 against both herpes- and poxviruses. Our results substantiate the potential therapeutic application of inhibiting BiP in cells infected by enveloped viruses from unrelated families. A primary concern when targeting host factors for therapeutic antiviral intervention is the potential for cytotoxicity. This concern is paramount when targeting BiP, which is critical for overall cell homeostasis ^13,35^. However, our results support the notion that BiP inhibition might be tolerable—we observed minimal cytotoxicity in three primary uninfected cell lines, including peripheral B-cells, lymphatic endothelial cells, and normal human dermal fibroblasts, at concentrations higher than those used to block viral replication. Moreover, *in vivo* studies have shown that BiP haploinsufficiency in aged mice had no significant adverse effects on body weight, organ integrity, behavior, memory, cancer, inflammation, or chemotoxic response ^67^. These observations and our results suggest that inhibiting BiP offers a promising therapeutic window for deploying broad-spectrum antivirals.

## Materials and Methods

### Cell Culture and Compounds

iSLK, iSLK.219, and normal human dermal fibroblasts (NHDFs/Lonza CC-2509) were grown in Dulbecco’s modified Eagle medium (DMEM; Invitrogen, Carlsbad, CA, USA) supplemented with 10% FBS, 200 µM of L-glutamine, and 100 U/mL of penicillin and streptomycin. iSLK.219 cells were maintained in 10 μg/mL of puromycin (Invivogen, San Diego, CA, USA). The Primary Effusion Lymphoma (PEL)-derived cells TREx-BCBL1-RTA (Jung Lab Lerner Research Institute at Cleveland Clinic), BC1 (CVCL_1079), and BC2 (CVCL_1856) (Manzano Lab, University of Arkansas for Medical Sciences) were grown in RPMI 1640 medium (Invitrogen, Carlsbad, CA, USA) supplemented with 10% fetal bovine serum (FBS; Invitrogen, Carlsbad, CA, USA), 200 μM of L-glutamine, and 100 U/mL of penicillin/streptomycin. Primary Dermal Lymphatic endothelial cells (LECs) from PromoCell (C-12217) were maintained in EBM-2 media (Lonza 00190860) supplemented with the EGM-2 MV bullet kit (CC-4147) at 37°C with a 5% CO2 atmosphere. Primary Peripheral B Cells (PPBSc, STEMCELL 70023) were thawed and maintained at 100,000 cells/mL in ImmunoCult™ Human B Cell Expansion media (STEMCELL 100-0645) at 37 degrees Celsius in 5% CO2. Cells were allowed to grow for 7-10 days before treatment with HA15.

The following drugs were used at the concentrations noted; Thapsigargin (Tg) (Tocris 1138) 100nM, Ceapin A7 (Sigma Aldrich SML2330) 6µM, HA15 (Selleckchem S8299) 1-50µM.

### Induction and assessment of KSHV reactivation and replication

Exogenous RTA expression was induced in iSLK.219 and TREx-BCBL-1-RTA cells by treatment with 1 μg/mL of doxycycline (Fisher Scientific, Waltham, MA, USA). To prevent viral DNA replication (Fig 1C), these cells were induced with Dox in the presence of phosphonoformate (PFA) 100µM (Sigma Aldrich P6801). Viral reactivation was evaluated by microscopy detection of the PAN-RFP reporter and immunoblot for viral proteins. To determine the efficiency of KSHV DNA replication, DNA was isolated from BCBL-1-RTA at the indicated times following reactivation using the Dneasy blood and tissue kit following manufacturer guidelines (Qiagen 69581). 20ng of total DNA was used for qPCR using primers for the KSHV gene ORF57: F: 5’GGGTGGTTTGATGAGAAGGACA3′ R: 5’CGCTACCAAATATGCCACCT, and Human Chromosome 11q (accession number AP002002.4) as a normalization control: F: 5′TAACTGGTCTTGACTAGGGTTTCAG3′ R: 5′ACCACAACAAAAAGCCTTATAGTGG3′

### Viruses

HSV1-US11-GFP (Patton strain) (Mohr lab, NYU School of Medicine) was propagated and titrated in Vero cells. HCMV-TB40/E-GFP (Murphy Lab SUNY) was propagated and titrated in NHDFs. Vaccinia Virus Western Reserve (ATCC VR-1354) was expanded in HeLa cells and titrated in BSC1 cells. KSHV.219 was generated from iSLK.219 cells treated with Dox 1ug/ml for 72h. The supernatant from lytic cells was collected, clarified, and filtered with a 0.45 um syringe filter. The virus was tittered by spinoculation (2000 rpm/2 h/Room temp) of uninfected iSLK in 6 well plates. Cells were incubated for 48 h following infection, trypsinized, and collected for flow cytometry in a Sony SH800 instrument. The percentage of cells expressing eGFP was determined by flow cytometry and used to calculate the number of fluorescence forming units (ffus) in each sample.

### Immunoblotting and Antibodies

Cells were washed and collected in 1X sample buffer (62.5 mM Tris-HCl (pH 6.8), 2% sodium dodecyl sulfate (SDS), 10% glycerol, 0.7 M b-mercaptoethanol). Samples were sonicated on ice to reduce viscosity. Cell lysates were fractionated by SDS-PAGE and transferred onto nitrocellulose membranes. Immunoblots were incubated with primary antibodies overnight at 4 °C, and immunoreactive bands were detected with HRP-conjugated secondary antibodies by enhanced chemiluminescence (ThermoFisher, Waltham, MA, USA) according to the manufacturer’s recommendations. All antibodies were used at a 1:1000 dilution in 3% BSA/1× TBST unless indicated. BiP (Cell Signaling Technologies 3117), Actin (1:30,000, Sigma Aldrich, St. Louis, MO, USA), GRP94 (Cell Signaling Technologies 2104), Calreticulin (Cell Signaling Technologies 2891, IRE1 (Cell Signaling Technologies 3294), IRE1-P Ser274 (Novus biotechnologies NB100-2323), XBP1s (Cell Signaling Technologies 40435), K8.1 (mAb clone 19B4), vIL6 (Advanced Biotechnology 13-214-050), KbZip (SCBT sc-69797), ORF45 (SCBT sc-53883), ORF57 (SBCT sc-135746). The LANA rabbit polyclonal antibody was raised against a synthetic peptide from the acidic domain of LANA (Polson and Ganem, unpublished). The antibody for K1 (1:100) was a generous gift from the Damania Lab at The University of North Carolina at Chapel Hill.

### CRISPRi-mediated knockdown

Synthetic DNA segments encoding the sgRNAs targeting ATF6 (5’GTTAATATCTGGGACGGCGG3′) or XBP1 (5’GCCGCCACGCTGGGAACCTA3′) were cloned into the BlpI and BstXI restriction sites in the pLG15 (CRISPRi) vector. The positive clones were confirmed by Sanger sequencing. Vesicular stomatitis virus (VSV) pseudotyped lentiviral production followed standard protocols. Briefly, 293METR packaging cell lines were transfected with the pLG15 lentiviral vector, VSV-G plasmid (pMD2.G Addgene 12259), and pCMV delta R8.2 (Addgene 12263). At 48h post-transfection, the viral supernatant was collected, clarified by centrifugation, and filtered through a 0.45 µm filter to remove cell debris. Viral particles were concentrated 5-fold using a regenerated cellulose centrifugal filter unit with a 100k MW cut-off (Amicon Ultracel 100k). The resulting lentivirus stock was used to transduce iSLK.219-dCas9-KRAB cells by spinoculation ^37^. Transduced iSLK.219 cells were maintained in 10 μg/mL of puromycin and were selected for BFP+/sgRNA+ expression by FACS in a Sony SH800 instrument. Knockdown of ATF6 and XBP1s was confirmed by qPCR or immunoblot, respectively.

### Reverse Transcription PCR (RT-PCR) and Quantitative PCR (qPCR)

Total cellular and viral RNA was isolated from cells using the RNAeasy Plus Mini kit (QIAGEN 74134) following manufacturers’ recommendations. Reverse-transcription (RT)-PCR was performed using 500-1000 ng of total RNA per RT reaction using the iScript Reverse Transcription Supermix. To remove excess genomic DNA, samples were subjected to Dnase (New England Biolabs Inc. M0303) treatment. PCR was done using as a template 1% of the resulting cDNA. For the detection of XBP1-s and XBP1-u mRNAs, we used the following primer pairs: XBP1u/s: F: 5’GGAGTTAAGACAGCGCTTGG3′ R: 5’ACTGGGTCCAAGTTGTCCAG3′. Products were separated on a 3% agarose gel and quantified by scanning densitometry (ImageJ). BiP mRNA abundance changes were measured by real-time RT-PCR analysis using the PowerUp SYBR Green Master Mix. All qPCR reactions were done in a C1000 Touch Thermal cycler with a CFX96 Real-Time System. Samples were normalized using 28S RNA. Primers: 28S: F: 5′AAACTCTGGTGGAGGTCCGT3′ R: 5′CTTACCAAAAGTGGCCCACTA3′, BiP (HSAP5): F: 5′AGTTCCAGCGTCTTTGGTTG3′ R: 5′TGCAGCAGGACATCAAGTTC3′

### siRNA-mediated knockdown

Small interfering RNAs targeting BiP (NM_005347) were ordered as a SMARTpool from Dharmacon (ON-TARGETplus Human HSPA5 siRNA L-008198-00-0005). The ON-TARGETplus Non-targeting Control Pool was used as a negative control (D-001810-10-05). iSLK.219 cells (2×10e5 cells/well) were transfected with 100nM of the siRNA mix using DharmaFect transfection reagent. At 24h post-silencing, cells were treated with 1 µg/mL Dox to induce viral lytic reactivation. BiP silencing was confirmed by immunoblot.

### RNA sequencing and analysis

Total cellular and viral RNA was isolated from iSLK.219 cells at 72h post reactivation in the presence or absence of 10µM HA15, using the RNAeasy Plus Mini kit (QIAGEN 74134) following manufacturers’ recommendations, including a DNAse treatment step. RNA sequencing libraries were generated using the NEBNext Ultra II RNA Library Prep Kit (New England BioLabs E7760) and sequenced using a 150bp paired-end protocol on an Illumina Novaseq 6000 instrument. Following demultiplexing, the sequenced reads were analyzed using the CZ ID platform (czid.org). Samples were aligned to the human genome GRcHg38, and all remaining reads were saved as non_host. These files were aligned to the KSHV genome GQ994935.1, and the transcripts were quantified using Salmon^68^. Heatmaps were generated and annotated in Prism.

### Fluorescence Assay

Primary normal human dermal (NHDF) cells were plated at a density of 30,000 cells per well in a 96-well plate. The following day, cells were pretreated for two hours with HA15 (DMSO final concentration 0.1%) and incubated at 37 °C. After pretreatment, NHDF cells were either mock-infected or infected with Herpes simplex virus 1 (HSV-1) US11-GFP (Patton strain) at MOI 0.01, Human cytomegalovirus (HCMV) EGFP (TB40/E strain) at MOI 0.1, or Vaccinia virus (VV) (Western Reserve strain) at MOI 0.1 and incubated for 1 h at 37 °C. After 1 h incubation, the supernatant from cells was removed and replaced with fresh DMEM and HA15. Cells were incubated at 37 °C for 24 h (HSV-1 and VV-infected cells) or 6 days (HCMV-infected cells). After respective incubation periods, the supernatant was removed and replaced with PBS. Because VV lacked a fluorescent reporter, infected cells were stained with primary antibody (Vaccinia Virus Polyclonal FITC Antibody ThermoFisher PA1-73191, 1:1000) and Hoechst 33342 (1:10,000). The fluorescent signal (GFP/Hoescht) was analyzed using the SpectraMax i3x plate reader. GFP fluorescence was measured at 485/535 and Hoescht fluorescence at 350/461.

### Cell Viability Assays

TREx-BCBL1-RTA, BC1, and BC2 cells were seeded at a density of 30,000 cells per well in a 96-well plate. The following day, cells were treated with increasing doses of HA15 (1µM, 5µM, 10µM, and 50µM). At 72h hours post-treatment, 10µl of the cells were stained with trypan blue and counted using the countess automated cell counter (ThermoFisher) to determine the number of live cells/ml and the percent cell death.

LEC viability was determined using the CellTiter-Glo Luminescent Cell Viability Assay (Promega). Uninfected and KSHV-infected LECs were seeded at a density of 7,500-10,000 cells per well of a white 96-well plate. The following day, cells were treated with HA15 (10µM). At 72h post-treatment, the media was replaced, and an equal volume of the CellTiter-Glo reagent was added to each well. The plate was incubated in the dark for 10-15 minutes before luminescence was read on a Victor^3^V 1420 Multilabel Counter (Perkin Elmer).

Primary normal human dermal (NHDF) cells were plated at a density of 30,000 cells per well in a 96-well plate. The following day, cells were treated with HA15 (DMSO final concentration 0.1%) and incubated at 37 °C. At 24 h or 6 days (corresponding to the viral infection period), 50 µL of supernatant was transferred to a new 96-well plate. 50 µL of CytoTox-ONE reagent was added to the plate and incubated at RT for 10 minutes. After 10 minutes, 25 µL of Stopping Reagent was added to the plate, and the plate was incubated at RT for 10 minutes. After incubation, the plate was transferred to the SpectraMax i3x plate reader, and fluorescence was read at 560/590 to determine percent cytotoxicity.

## Acknowledgments

We thank the CZ BioHub genomics team for their help with RNA sequencing. We are grateful for the support, scientific discussions, and critical proofreading from S. Schmid, A. Kistler, and R. Aviner at CZ BioHub, San Francisco; D. Acosta-Alvear, F. Braig-Karzig and F. Zappa at Altos Labs; and Z. Aralis and D. Proctor at UCSB. This work was funded by a grant to CS by the UC Research Initiatives, Cancer Research Coordinating Committee, and an NSF Bridge to Doctorate fellowship to GN.

## Supplementary Figures

**Supplementary Figure 1. KSHV reactivation in iSLK.219 cells follows a cascade of gene expression.**

Latently infected iSLK.219 cells were induced to enter the lytic cycle by exogenous expression of RTA following Dox (1µg/ml) treatment. **(A.)** Imaging of cells at 72h post reactivation showing the expression of the lytic PAN-RFP marker in the population. **(B.)** Immunoblot for viral proteins in iSLK.219 lysates collected at the indicated time points. Images are representative of 3 independent biological replicates. Actin: loading control.

**Supplementary Figure 2. BiP is post-transcriptionally upregulated independently of ATF6 and XBP1.**

**(A-C.)** IRE1 is phosphorylated in lytic iSLK.219 cells without detectable XBP1 splicing. Cells were reactivated by treatment with Dox (1µg/ml). At the indicated times, the cells were treated with Tg (100nM) for 4h to induce acute ER stress. **(A.)** Whole-cell lysates were collected and analyzed by immunoblot for total (IRE1) or phosphorylated IRE1 (IRE1-P), spliced XBP1 (XBP1s), and actin (loading control). **(B.)** RT-PCR detection of unspliced (u) and spliced (s) XBP1 mRNA. **(C.)** Image densitometry quantification of the data in (B). **(D-H.)** XBP1 and ATF6 are not required for BiP protein upregulation or infectious virus production during the KSHV lytic cycle. **(D.)** CRISPRi-based knockdown of XBP1 (XBP1-KD) in iSLK.219-dCas9 cells. Cells (NS and XBP1-KD) were induced with Dox (1µg/ml) for 24h. Cells were treated with Tg (100nM) for 4h before collection. (left) Whole cell lysates were analyzed by immunoblot. Actin: loading control. **(E.)** The supernatants of cells treated as in (D) were collected and used to spinoculate uninfected iSLK cells. The percent of GFP expression was determined by automated cell counting and used as a proxy for infectious virus levels in the supernatants. **(F.)** iSLK.219 cells were treated with the ATF6 inhibitor CeapinA7 (6µM) for 2h before induction with Dox (1µg/ml). Whole-cell lysates were collected at the indicated times and analyzed by immunoblot. **(G.)** CRISPRi-based knockdown of ATF6 (ATF6KD) in iSLK.219-dCas9 cells. ATF6-KD cells were treated as in (D). **(H.)** Supernatants from ATF6-KD cells were collected and processed as described in (E). *N*=3 independent biological replicates. Values in (C, E, F) are average ±SEM. Statistical significance was calculated using a one-way ANOVA (*P=0.01) in (C) or a paired t-test (E and H).

**Supplementary Figure 3: BiP levels do not increase during the KSHV lytic cycle in TREx-BCBL-1 cells.**

TREx-BCBL-1 cells were reactivated with Dox (2 µg/ml). At 4h before collection, cells were treated with Tg (100nM) for 4h to induce acute ER stress. Whole-cell lysates were collected at the indicated times. Actin: loading control.

**Supplementary Figure 4: HA15 treatment of iSLK.219 does not induce XBP1s expression.**

Latent iSLK.219 cells were reactivated in the presence or absence of HA15 (10µM). Whole-cell lysates collected at the indicated times were analyzed by immunoblot using an antibody specific for XBP1s. Actin: loading control.

**Supplementary Figure 5. HA15 has a cytostatic effect on PEL-derived cells.**

**(A-D)** HA15 treatment causes cytostasis in BC-1 and BC-2 cells latently co-infected with KSHV and EBV. Cells were treated with increasing doses of HA15 (0-50 μM) for 72h. The total number of viable **(A.)** and the percent of dead BC-1 cells **(B.)** were determined by automated cell counting following trypan blue staining. The total number of viable **(C.)** and the percent of dead BC-2 cells **(D.)** were determined by automated cell counting following trypan blue staining. *N*=3 independent biological replicates. Values are average ±SEM.

## Notes

### Competing Interest Statement

The authors have declared no competing interest.

