## Supplementary figures for "BiP/GRP78 is a pro-viral factor for diverse dsDNA viruses that promotes the survival and proliferation of cells upon KSHV infection"

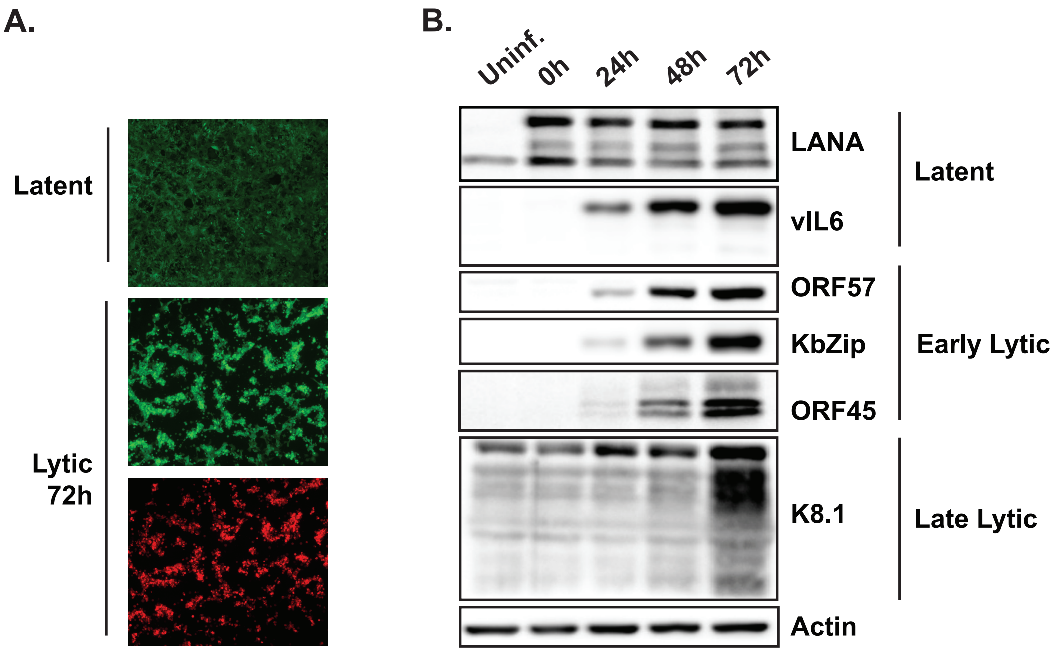


**Supplementary Figure 1. KSHV reactivation in iSLK.219 cells follows a cascade of gene expression.**

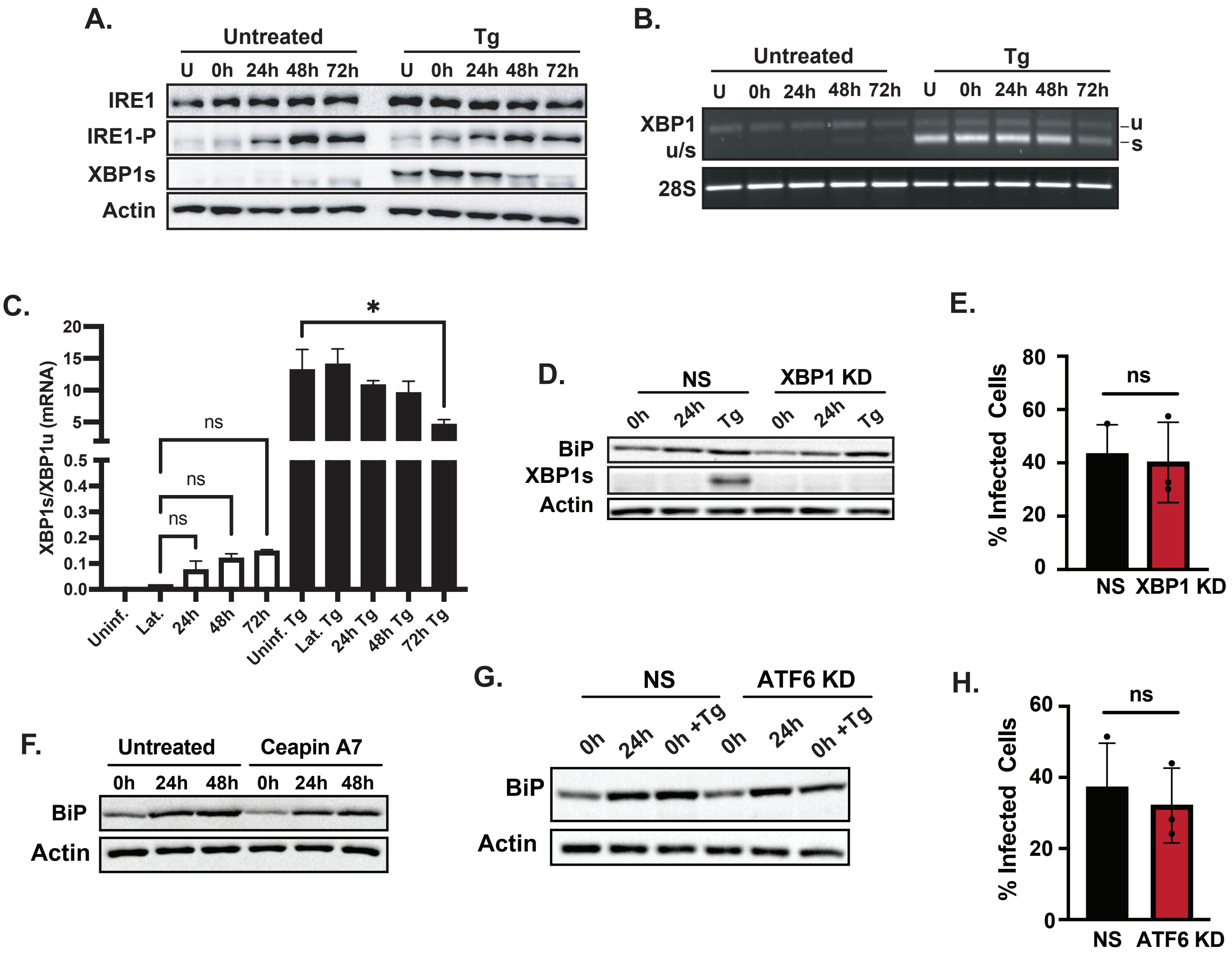


**Supplementary Figure 2. BiP is post-transcriptionally upregulated independently of ATF6 and XBP1.**

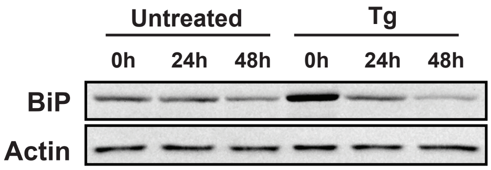


**Supplementary Figure 3: BiP levels do not increase during the KSHV lytic cycle in TREx-BCBL-1 cells.**

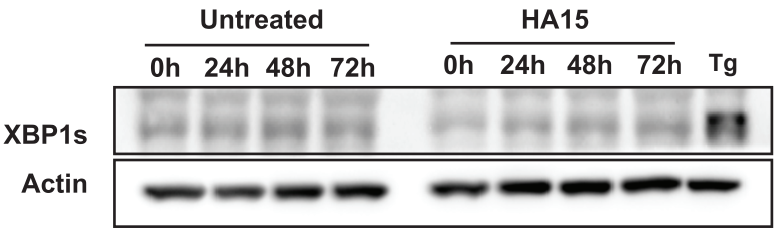


**Supplementary Figure 4: HA15 treatment of iSLK.219 does not induce XBP1s expression.**

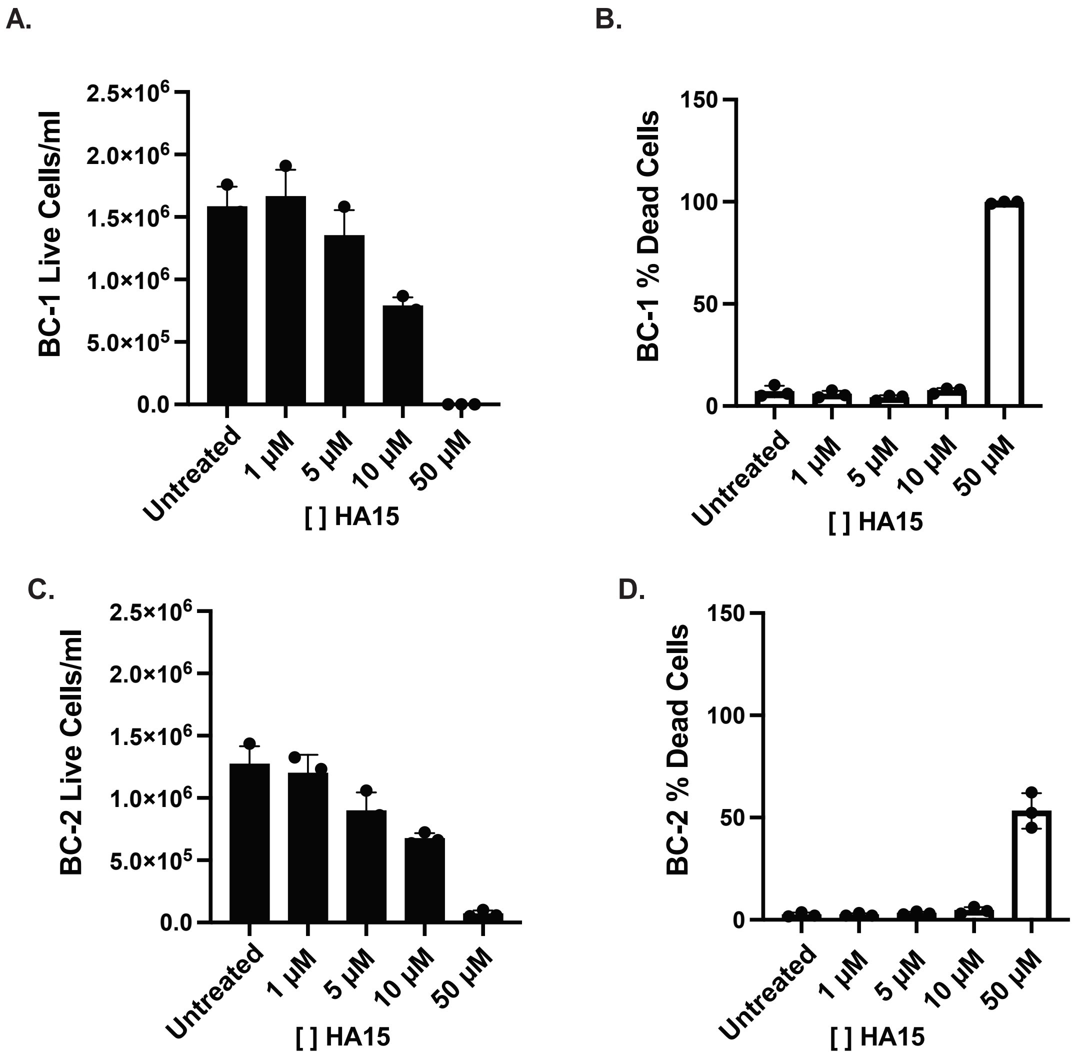


**Supplementary Figure 5. HA15 has a cytostatic effect on PEL-derived cells.**
